## Supplementary material for "Sex-dependent effects of *Setd1a* haploinsufficiency on development and adult behaviour": Supplemntary Information

**Supplementary Information**

File contains:


| Page | Item |
|---|---|
| 2 | Supplementary Methods Figure 1: Confirmation of *Setd1a* haploinsufficiency in the *Setd1a*$^{+/-}$ model. |
| 3 | Supplementary Methods Table 1: Samples sizes for evaluation of embryo and placenta size. |
| 4 | Supplementary Methods Figure 2: Methods for RNAseq |
| 5 | Supplementary Methods Figure 3: Pilot studies for pharmacological investigations |
| 6 | Supplementary Results Figure 1: Trajectory of Setd1a expression across neurodevelopment in WT mice and confirmation of *Setd1a* haploinsufficiency in the *Setd1a*$^{+/-}$ model. |
| 7 | Supplementary Results Table 1: Details of the litters, genotype and sex ratios of the samples used in the pre- and postnatal assessments. |
| 7 | Supplementary Results Figure 2: Acoustic startle response analysis per trial |
| 8 | Supplementary Methods Figure 3: Replication of acoustic startle and prepulse inhibition effects (Main text Fig. 4a and 4b) in a separate cohort of *Setd1a*$^{+/-}$ and WT mice. |
| 8 | Supplementary Results Table 2: Supporting data for the novel object recognition test. |

**Supplementary Methods Figure 1: Confirmation of *Setd1a* haploinsufficiency in the *Setd1a*$^{+/-}$ model.**

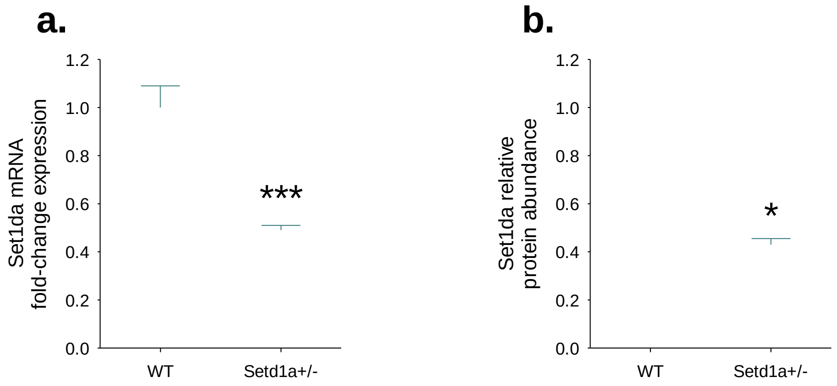


To confirm *Setd1a* haploinsufficiency in our model, levels of *Setd1a* mRNA and protein were quantified in whole brains dissected at E13.5 N=8 for WT and *Setd1a*$^{+/-}$ tissue). Haploinsufficiency was confirmed in the *Setd1a*$^{+/-}$ model, by demonstrating that *Setd1a* mRNA expression was reduced by 48.8% at E13.5 in *Setd1a*$^{+/-}$ brain compared to WT (Fig. 1c, $t_{14}$=9.18, p<0.001), with comparable reductions in levels of SETD1A protein (46.3 % reduction) (Fig. 1d, $t_8$=2.71, p=0.03). This magnitude of reduction is consistent with and confirmed that *Setd1a* knockdown to half of WT levels was successfully achieved. * and *** shows significant comparison to WT at p<0.05 and p<0.001, respectively. Data shows mean±SEM.

Methods: RNA extraction was performed using a Direct-zol™ RNA Miniprep Kit (Zymo, UK). 1 µg total RNA was used for cDNA synthesis using RNA to cDNA EcoDry™ Premix (double-primed) kits (Clontech, UK). qRT-PCR reactions were performed in triplicate using a Corbett Rotorgene 6000 Real-Time PCR machine with Sensimix SYBR No-Rox (Bioline, UK) and intron-spanning primers.

| Target | Forward primer | Reverse primer | Product size (bp) |
|---|---|---|---|
| *Setd1a* | CCCTCCCGGTTCCTAAGTTT | CATTGTCATTGAGCCTCGCA | 90 |
| *Hprt* | GCGATGATGAACCAGGTTATGA | GCCTCCCATCTCCTTCATGA | 146 |
| *Dynein* | GACCTCAGGCTCAGACGAAGAC | AAGACGCTCATGGCATCACA | 116 |
| *B2m* | TTCTGGTGCTTGTCTCACTGA | CAGTATGTTCGGCTTCCCATTC | 104 |

The geometric mean of Ct values across three housekeeping genes (*Hprt*, *Dynein*, and *B2m*) were used as endogenous controls to normalise *Setd1a* expression levels using the ΔΔCT method[25]. Protein was extracted from brain homogenates in RIPA buffer (Sigma, UK) containing cOmplete™ Mini Protease Inhibitor Cocktail (Roche, Switzerland). A Pierce™ BCA Protein Assay kit (Thermo Scientific, UK) was used to quantify protein concentration. Samples were diluted in protein loading buffer (LI-COR, UK) containing 0.05 % (v/v) 2-Mercaptoethanol (Sigma, UK) and denatured by heating at 95 °C for five minutes. 20 µg total protein per sample was separated by SDS-PAGE using a NuPAGE™ 4-12 % Tris-Acetate gel (Invitrogen, UK) and NuPAGE™ Tris-Acetate SDS Running Buffer (Invitrogen, UK). Proteins were transferred to a 0.45 µm pore size nitrocellulose membrane (Invitrogen, UK) in NuPAGE™ Transfer Buffer (Invitrogen, UK) containing 10 % (v/v) methanol (Fisher Scientific, UK). To enable normalisation of *SETD1A* protein abundance, membranes were stained for total protein using REVERT™ Total Protein Stain (LI-COR, UK). Odyssey TBS Blocking Buffer (LI-COR, UK) was used to block membranes for one hour at room temperature. Membranes were incubated overnight at 4 °C with 1:1,000 polyclonal *Setd1a* antibody (Bethyl Laboratories, USA). TBS-T (1M NaCl, 1M Tris-HCl, 0.2 % (v/v) Tween 20) was used to wash the membrane four times (5 minutes per wash) before incubation in 1:10,000 IRDye 800CW goat anti-rabbit secondary antibody (LI-COR, UK) for one hour at room temperature. Wash steps were repeated prior to imaging using an Odyssey CLx and protein quantification using Image Studio software (LI-COR, UK).

**Supplementary Methods Table 1: Samples sizes for evaluation of embryo and placenta size**

| | Number of litters | WT Male | *Setd1a*$^{+/-}$ Male | WT Female | *Setd1a*$^{+/-}$ Female | Total |
|---|---|---|---|---|---|---|
| **E11.5** | 4 | 11 | 5 | 6 | 6 | 28 |
| **E13.5** | 5 | 13 | 11 | 14 | 8 | 46 |
| **E18.5** | 6 | 10 | 11 | 10 | 13 | 44 |

### Supplementary Methods Figure 2: Methods for RNAseq

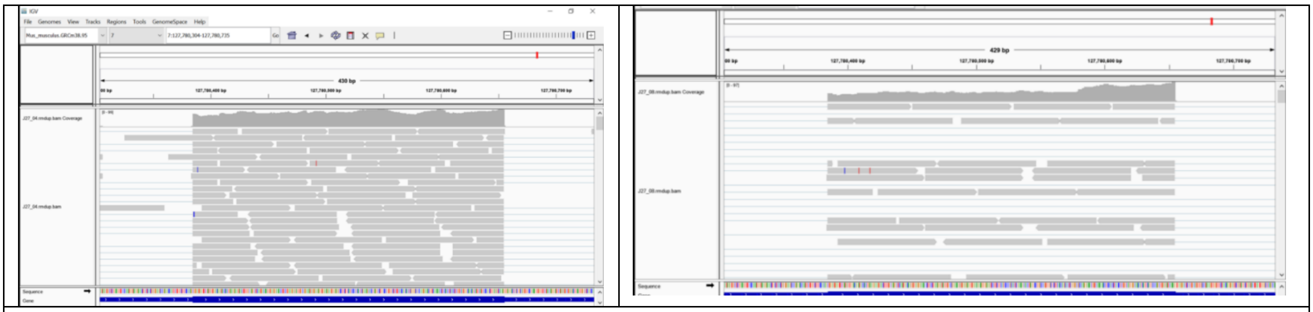

RNA purity was assessed using a NanoDrop 8000 spectrophotometer (Thermo Fisher Scientific, UK). RNA concentration was quantified using a QubitTM RNA High Sensitivity Assay Kit and Qubit 2.0 Fluorometer (Invitrogen, UK). RNA integrity was assessed using a Bioanalyzer RNA 6000 Nano Assay (Agilent, UK) and 2100 Bioanalyzer system (Agilent, UK). All samples had an RNA Integrity Number of at least 9.7. Library preparations were performed using a KAPA mRNA Hyperprep kit (Roche, Switzerland) according to manufacturer's instructions, with 1 µg total RNA as input. Following library amplification, fragment size (mean = 374.9, SD = 12.5) was determined using a High Sensitivity DNA kit (Agilent, UK). A QubitTM dsDNA High Sensitivity Assay Kit (Invitrogen, UK) was used to determine library concentration. Reads were trimmed to remove adapters and low-quality bases using Trimmomatic[1] with default parameters. Reads were mapped to the mouse reference genome (GRCm38) using STAR[2]. The mean number of reads mapped was 97.2% (SD = 0.5%). Read counts were generated using featureCounts[3] to allocate reads to genomic features using the mouse Ensembl gene annotation (GRCm.38.95). Coverage at exon 4 of *Setd1a* was substantially reduced in *Setd1a*+/- E13.5 brain, indicating that recombination had occurred (see figure, screenshot from Integrative Genomics Viewer showing substantially more reads aligning to exon 4 in WT (left) compared to *Setd1a*$^{+/-}$ (right) mice). See main text for other details.


1. Bolger, A. M., Lohse, M., & Usadel, B. (2014). Trimmomatic: a flexible trimmer for Illumina sequence data. *Bioinformatics*, *30*(15), 2114–2120. doi: 10.1093/bioinformatics/btu170
2. Dobin, A., Davis, C. A., Schlesinger, F., Drenkow, J., Zaleski, C., Jha, S., … Gingeras, T. R. (2013). STAR: ultrafast universal RNA-seq aligner. *Bioinformatics*, *29*(1), 15–21. doi: 10.1093/bioinformatics/bts635
3. Liao, Y., Smyth, G. K., & Shi, W. (2014). featureCounts: an efficient general purpose program for assigning sequence reads to genomic features. *Bioinformatics*, *30*(7), 923–930. doi: 10.1093/bioinformatics/btt656

### Supplementary Methods Figure 3: Pilot studies for pharmacological investigations

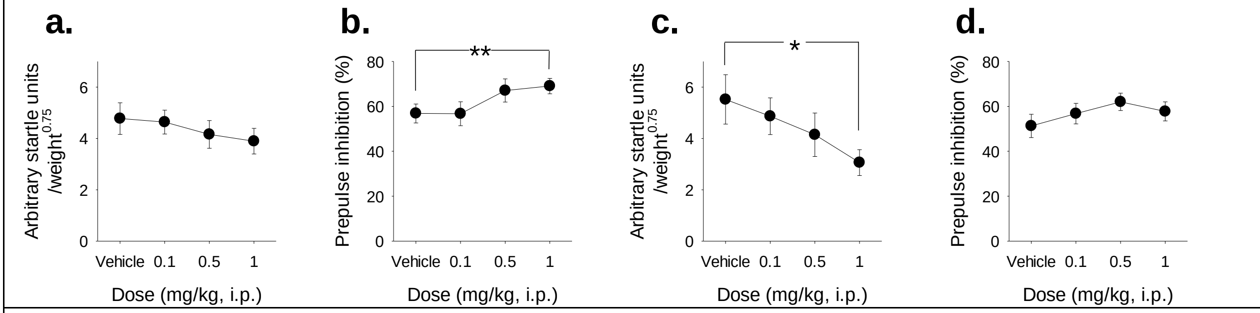


Previous work in rodent models has shown that PPI deficits induced by D2 receptor agonists are reversed by both haloperidol and risperidone, whereas NMDAR antagonist induced deficits are generally insensitive to haloperidol but can be reversed by risperidone[1,2], therefore we used these drugs to gauge their effects on *Setd1a*$^{+/-}$ mice. Using dose ranges, and a 30 min pre-test delay, from this literature with an aim to test effects of haloperidol and risperidone in WT mice to identify an appropriate dose range for the main experiment. Two groups of 3 months old male C57BL/6J mice (N=17 and N=16, for the haloperidol and risperidone studies, respectively) were used, using a protocol consisting of 25 trials, with 120 dB pulse-alone startle stimuli and 8 and 16dB prepulse trials (see main text for further description) (for analysis, PPI was pooled across 8 and 16 dB above background prepulses). Haloperidol did not significantly affect acoustic startle responding (a, main effect of DOSE, $F_{3,48}$=1.43, p=0.25), however the 1.0mg/kg dose significantly increased PPI relative to vehicle (b, main effect of DOSE, $F_{3,48}$=4.11, p=0.01), confirmed by *post hoc* testing (p=0.04 for the comparison between vehicle and 1.0mg/kg dose, other comparisons were n.s.). Administration of risperidone significantly reduced the startle response (c, main effect of DOSE, $F_{3,45}$=6.42, p=0.001). *Post-hoc* tests revealed a significantly reduced startle response at 1.0 mg/kg (p=0.03) but not with the other doses of risperidone. Risperidone did not affect PPI at any dose (main effect of DOSE, $F_{3,45}$=2.12, p=0.15). These results showed that 0.5 mg/kg was the highest dose of haloperidol and risperidone tested that did not have significant effects on basal sensorimotor gating in WT mice. Thus, in the main experiment, all mice were tested at this dosage in the first instance to explore drug effects on sensorimotor gating in *Setd1a*+/- mice in the absence of non-specific effects. * and ** shows significant main effect of DOSE at p<0.05 and p<0.01, respectively Data shows mean±SEM

1. Geyer, M. A., Krebs-Thomson, K., Braff, D. L., & Swerdlow, N. R. (2001). Pharmacological studies of prepulse inhibition models of sensorimotor gating deficits in schizophrenia: a decade in review. *Psychopharmacology*, *156*(2–3), 117–154. doi: 10.1007/s002130100811
2. Swerdlow, N. R., Talledo, J., Sutherland, A. N., Nagy, D., & Shoemaker, J. M. (2006). Antipsychotic Effects on Prepulse Inhibition in Normal 'Low Gating' Humans and Rats. *Neuropsychopharmacology*, *31*(9), 2011–2021. doi: 10.1038/sj.npp.1301043

**Supplementary Results Figure 1: Trajectory of Setd1a expression across neurodevelopment in WT mice.**

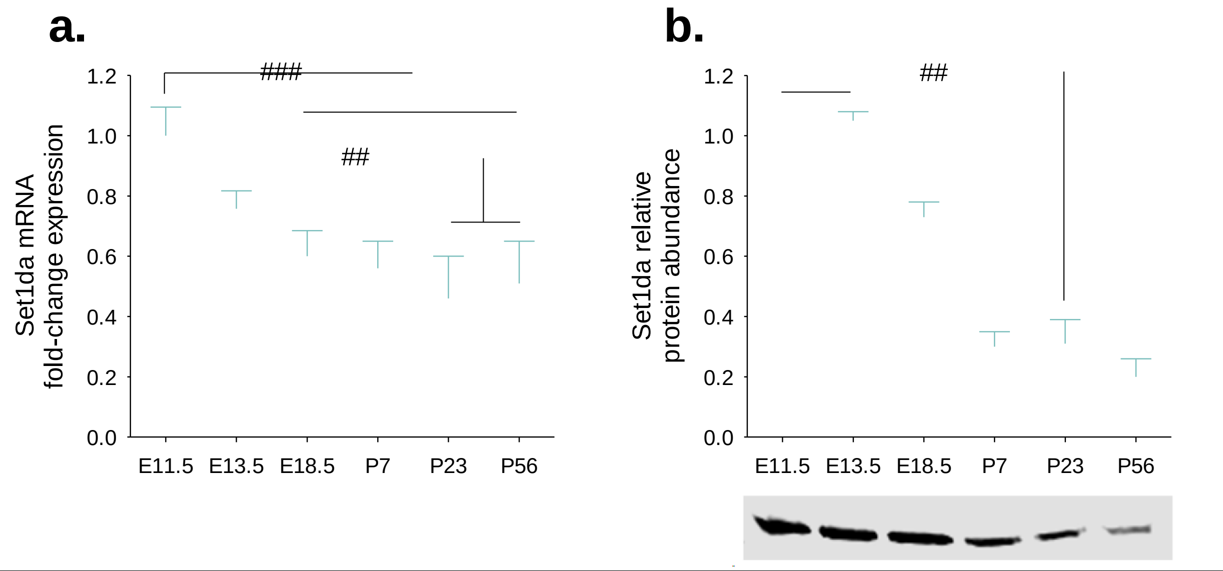


To determine the trajectory of Setd1a expression across neurodevelopment whole brains (E11.5, E13.5 and E18.5) or hemibrains (P7, P23 and P56), from at least two separate litters per timepoint, were obtained. Timed-matings were conducted between C57BL/6J x C57BL/6NTac parents and pregnant dams were culled by cervical dislocation. Sample analysis followed the methods detailed in Supplementary Methods Figure 1. Setd1a expression significantly reduced with age (a, main effect of AGE, $F_{5,30}=11.65$, $p<0.001$) in the brains of WT mice. *Post-hoc* tests showed that Setd1a expression levels were significantly higher at E11.5 than other timepoints (all $p<0.002$), except E13.5 ($p=0.26$). Expression levels at E13.5 were not significantly different from E18.5 ($p=0.32$) or P7 ($p=0.08$) but were significantly higher than at P23 ($p=0.003$) and P56 ($p=0.02$). There was no further significant decline in Setd1a expression levels after E18.5 (all $p>0.33$). Similar results were obtained at the protein level (b) with a significant main effect of AGE ($F_{5,24}=6.78$, $p<0.001$), which reflected significantly increased Setd1a expression at E11.5 and E13.5 relative to postnatal timepoints (all $p<0.02$). No significant difference was observed between E11.5 and E13.5 or E18.5 (both $p>0.60$). Protein levels did not change after E18.5 (all $p>0.17$). Together, these data suggest that Setd1a expression levels are higher in the developing brain during (mid) gestation, followed by down-regulation in the postnatal brain. ## and ###, signify significant main effects of AGE at $p<0.01$ and $P<0.001$, respectively. Data shows mean±SEM.

**Supplementary Results Table 1: Details of the litters, genotype and sex ratios of the samples used in the pre- and post-natal assessments.**

| | Number of litters | Median litter size | WT:Setd1a$^{+/-}$ | | Males:Females | |
|---|---|---|---|---|---|---|
| | | | (Mean±SEM) | Chi$^2$ test | (Mean±SEM) | Chi$^2$ test |
| E11.5 | 4 | 7 | 0.39±0.09 | $X^2_2$=0.50, p>0.05 | 0.57±0.05 | $X^2_3$=0.00, p>0.05 |
| E13.5 | 5 | 10 | 0.43±0.08 | $X^2_3$=0.00, p>0.05 | 0.53±0.03 | $X^2_4$=0.60, p>0.05 |
| E18.5 | 6 | 7 | 0.55±0.07 | $X^2_4$=0.67, p>0.05 | 0.49±0.12 | $X^2_5$=0.00, p>0.05 |
| Postnatal (P28) | 13 | 7 | 0.45±0.07 | $X^2_{10}$=1.39, p>0.05 | 0.44±0.07 | $X^2_{10}$=1.39, p>0.05 |

Note: WT: Setd1a$^{+/-}$ ratio is the of number of Setd1a$^{+/-}$ samples/total samples and M:F ratio is calculated as number of male samples/total samples.

**Supplementary Results Figure 2: Supplementary Results Figure 2: Acoustic startle response analysis per trial.**

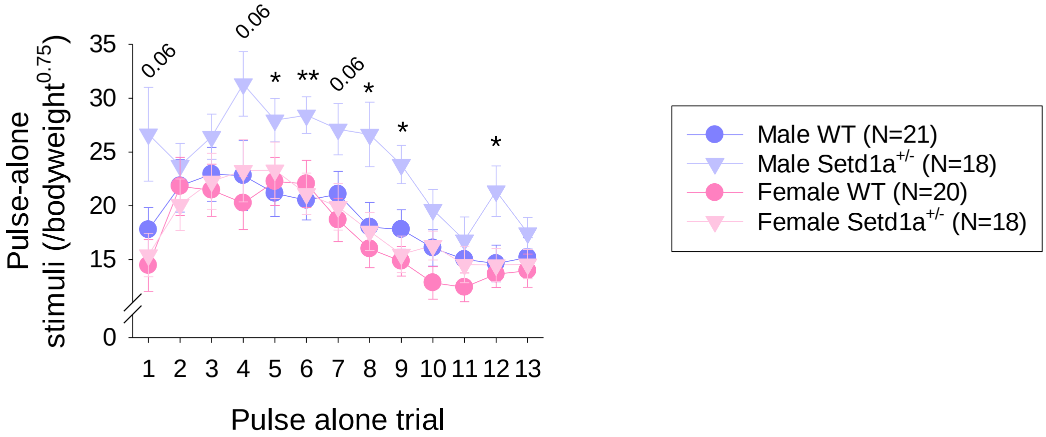


Analysis of the individual pulse-alone trials demonstrate that male Setd1a$^{+/-}$ mice showed elevated startle responding across the majority of trials as represented by significant main effects of GENOTYPE ($F_{1,72}$=4.51, p=0.04) and SEX ($F_{1,72}$=6.29, p=0.01). Simple effects analysis shows that in the male data the effect of GENOTYPE was significant ($F_{1,37}$=6.29, p=0.02), whereas there were no differences between female WT and Setd1a$^{+/-}$ mice ($F_{1,35}$=0.18, p=0.67) and all individual comparisons were non-significant (p>0.05). *post hoc* comparisons between male WT and Setd1a$^{+/-}$ mice, indicated on the graph, show that male Setd1a$^{+/-}$ mice show an increased response to the first startle stimulus, and then hyperstartle and demonstrate reduced habituation relative to male WT, and female mice through the remainder of the session. Note: mean of trials 4 to 13 is presented in the main paper figure 3a. * and **, signify significant main effects of GENOTYPE at p<0.05 and P<0.01 for comparison of male mice, respectively. Data shows mean±SEM.

**Supplementary Results Figure 3: Replication of acoustic startle and prepulse inhibition effects (Main text Fig. 3a and 3b) in a separate cohort of *Setd1a*$^{+/-}$ and WT mice.**

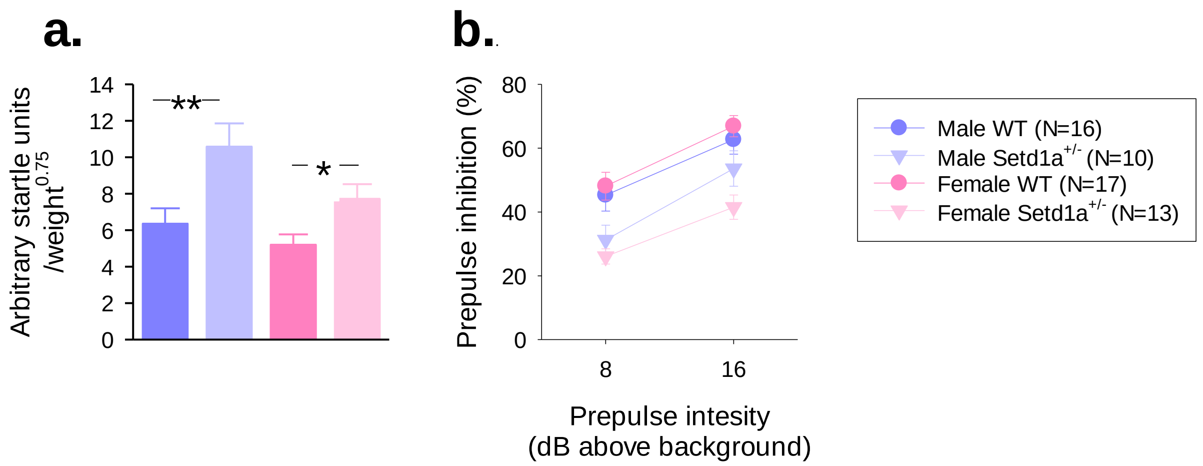


Consistent with the previous results (Main text Fig. 3a and 3b), there was a significant main effect of GENOTYPE on startle responding (a., $F_{1,52}$=15.58, p<0.001) which further confirmed that startle magnitude was significantly greater in *Setd1a*$^{+/-}$ mice than their WT littermates. The difference between male WT and *Setd1a*$^{+/-}$ mice was greater than before ($t_{24}$=2.89, p=0.008), and we also also observed a significant tendency for female *Setd1a*$^{+/-}$ mice to have elevated startled responses in this replication study ($t_{28}$=2.61, p=0.014). There was no GENOTYPE*SEX interaction ($F_{1,52}$=1.01, p=0.32), as before. For prepulse inhibition, there was a significant main effect of GENOTYPE ($F_{1,52}$=19.79, p<0.001) replicating the reduced PPI shown by *Setd1a*$^{+/-}$ mice (b). The interaction between GENOTYPE*PPI-INTENSITY was not significant ($F_{1,52}$=0.45, p=0.83) showing that this attenuation in PPI was consistent at both prepulse intensities used, even though the 16 dB prepulse induced greater PPI (main effect of PPI-INTENSITY, $F_{1,52}$=94.76, p<0.001). Thus, these data replicate the initial findings and demonstrate that increased ASR (particularly in males) and decreased PPI is a robust phenotype of *Setd1a* haplosufficiency. * and ** shows significant main effect of GENOTYPE at p<0.05 and p<0.01, respectively. Data shows mean±SEM.

**Supplementary Results Table 2: Supporting data for the novel object recognition test.** Mean (SD) of acquisition time (time taken to achieve 40 seconds of object exploration) and the individual object exploration times at test for the 30 minute and 24 hour retention intervals.

| | **30 mins** | | **24 hours** | |
|---|---|---|---|---|
| **Dependent variable** | **WT** | **KO** | **WT** | **KO** |
| Acquisition time (mins) | 6.9 (3.8) | 7.3 (4.2) | 7.1 (4.0) | 6.9 (4.1) |
| Novel object exploration time (s) | 19.2 (9.0) | 20.1 (11.0) | 8.1 (5.2) | 8.0 (5.4) |
| Familiar object exploration time (s) | 10.9 (8.7) | 9.5 (6.8) | 5.7 (4.0) | 6.1 (3.6) |

**End of document**
